# ADAMDEC1 maintains a novel growth factor signaling loop in cancer stem cells

**DOI:** 10.1101/531509

**Authors:** James S. Hale, Ana Jimenez-Pascual, Anja Kordowski, Jamie Pugh, Shilpa Rao, Daniel J. Silver, Tyler Alban, Defne Bayik Watson, Rui Chen, Thomas M. McIntyre, Giorgio Colombo, Giulia Taraboletti, Karl O. Holmberg, Karin Forsberg-Nilsson, Justin D. Lathia, Florian A. Siebzehnrubl

## Abstract

Glioblastomas (GBM) are lethal brain tumors where poor outcome is attributed to cellular heterogeneity, therapeutic resistance, and a highly infiltrative nature. These characteristics are preferentially linked to GBM cancer stem cells (GSCs), but how GSCs maintain their stemness is incompletely understood and the subject of intense investigation. Here, we identify a novel signaling loop that induces and maintains GSCs. This loop consists of an atypical metalloproteinase, a disintegrin and metalloproteinase domain-like protein decysin 1 (ADAMDEC1), secreted by GSCs. ADAMDEC1 rapidly solubilizes fibroblast growth factor-2 (FGF2) to stimulate FGF receptor 1 (FGFR1) expressed on GSCs. This signaling axis induces upregulation of Zinc finger E-box-binding homeobox 1 (ZEB1) that regulates ADAMDEC1 expression, creating a positive feedback loop. Genetic or pharmacological targeting of components of this axis attenuates self-renewal and tumor growth. These findings reveal a new signaling axis for GSC maintenance and highlight ADAMDEC1 and FGFR1 as potential therapeutic targets in GBM.

**Statement of Significance:** Cancer stem cells (CSC) drive tumor growth in many cancers including glioblastoma. We identified a novel sheddase, a disintegrin and metalloproteinase domain-like protein decysin 1, that initiates a fibroblast growth factor autocrine loop to promote stemness in CSCs. This loop can be targeted to reduce glioblastoma growth.

## Introduction

Glioblastoma (GBM) is a uniformly fatal disease with a median survival of approximately 20 months after diagnosis (1–3). GBM represents a prototypical example of a highly heterogeneous tumor with multiple distinct, identifiable molecular subclasses within a single tumor (4, 5). Frequent tumor recurrence and poor outcome are thought to be a consequence of resident GBM cancer stem cell (GSC) populations resistant to current therapies (6–8). Thus, GSCs are a candidate population for tumor recurrence (9–12). A recent study demonstrated that GBM contains GSC populations that produced non-stem cancer cell (NSCC) progenies, with only GSCs capable of propagating tumor formation (13). How GSCs are maintained across the changing tumor microenvironment within hypoxic, vascular, or invasive niches (14) remains unclear.

Tumor progression is frequently linked to the secretion of metalloproteinases that enable tissue invasion and intravasation by cancer cells via extracellular matrix degradation. This also causes a release of trophic factors to stimulate tumor growth, dispersal, and modulation of inflammatory responses. Members of the A Disintegrin and Metalloproteinase (ADAM) family of zinc-dependent proteinases contribute to GBM therapeutic resistance and invasiveness (15, 16), as well as to the regulation of GSCs (17–19). ADAMDEC1 is a soluble member of this family that is novel in mammals. It has restricted hydrolytic capacity due to a substitution in an active site residue (20) and selectively solubilizes growth factors from immobile precursor forms (21). Whether ADAMDEC1 solubilizes additional ligands, and whether this contributes to GBM growth and progression, has yet to be determined.

Trophic factors from the tumor microenvironment are essential to GBM growth and GSC maintenance. Fibroblast growth factor 2 (FGF2) is crucial in normal neural development and stem cell function, and a known oncogenic factor in GBM (22). FGF2 promotes glioma growth and vascularization (23) and GSC self-renewal (24). Nevertheless, how FGF2 specifically contributes to GSC functions is incompletely understood.

We now find ADAMDEC1 initiates FGF2 signaling through FGFR1 and mediates GSC self-renewal and maintenance through induction of the stem cell transcription factor, ZEB1. ZEB1 additionally induces expression of ADAMDEC1, completing a positive feedback loop and contributing to GSC maintenance in the tumor microenvironment. Importantly, we show that pharmacological intervention at the level of FGF2 can disrupt this loop and may constitute a translational therapeutic strategy.

## Results

Given the role for some of the ADAM family of metalloproteinases in GBM growth and progression (17, 19), we assessed the entire family for other members that are elevated in GBM and may contribute to tumor growth. We interrogated the bioinformatics database GlioVis (25) for mRNA expression levels of ADAM family members across multiple GBM datasets (Fig. 1A). When normalized to brain expression levels, ADAMDEC1 emerged among the top proteases upregulated in GBM. Further investigation of these candidates using TCGA and LeeY data sets revealed that ADAMDEC1 was the only protease where increased expression associated with poorer prognosis of GBM patients (Fig. 1B, S1). TCGA data also demonstrates that ADAMDEC1 mRNA levels increase with glioma grade (Fig. 1C). We performed immunohistochemistry on patient specimens to test whether this extends to the protein level and found elevated ADAMDEC1 in GBM (Fig. 1D). Fluorescence immunostaining in patient-derived xenograft models revealed that ADAMDEC1 was expressed by tumor cells, and not host microglia (Fig. 1E).

**Figure 1:**
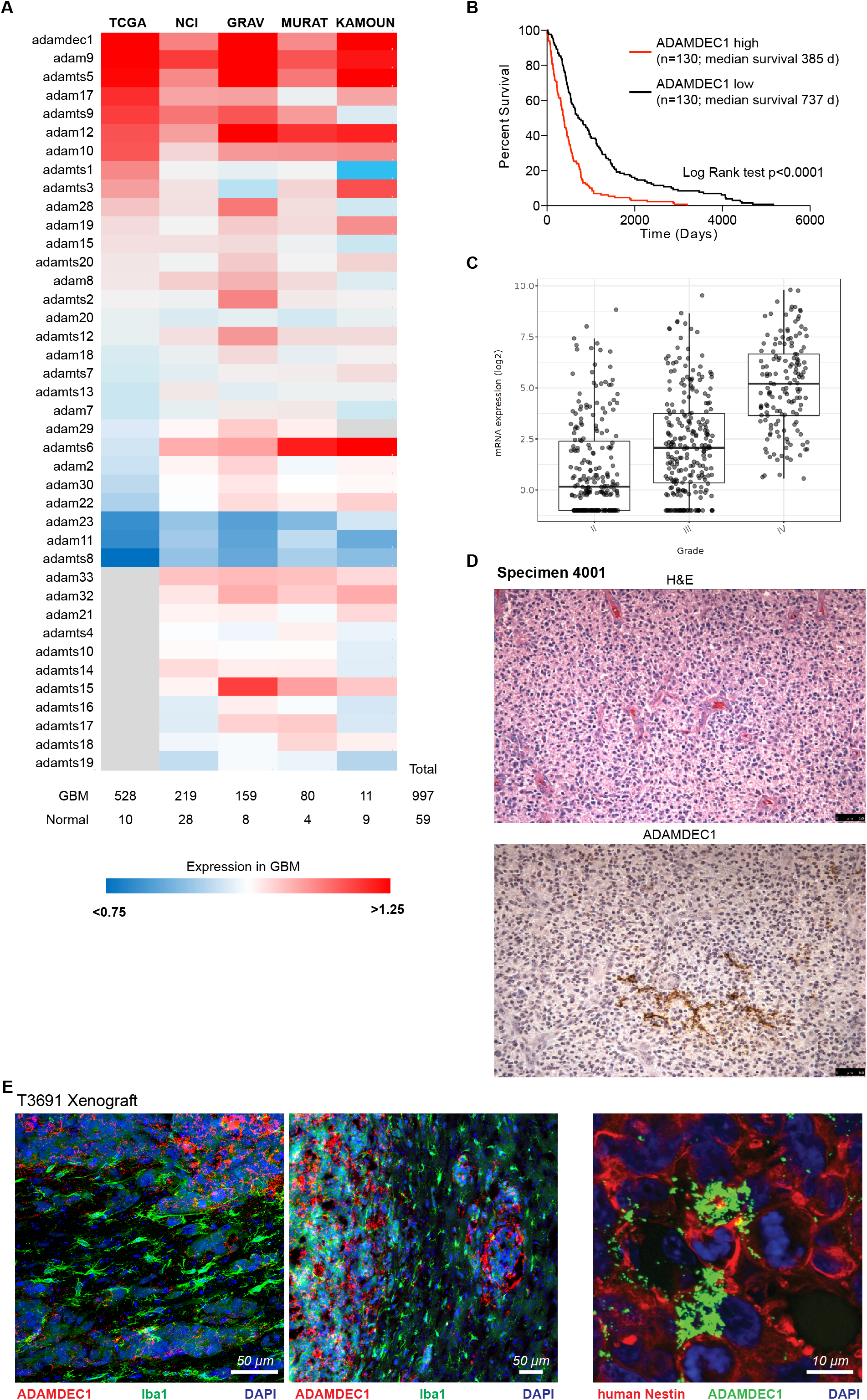
ADAMDEC1 is associated with malignancy in GBM. **(A)** Expression of ADAM family metalloproteinases across different GBM datasets. Expression levels were normalized to normal brain tissue samples and are presented as fold change. ADAMDEC1 is highly expressed across all datasets. **(B)** Kaplan-Meier survival analysis of TCGA data, stratified for above-median (high) or below-median (low) gene expression shows a significantly poorer survival in ADAMDEC1 high tumors. **(C)** ADAMDEC1 mRNA expression levels are significantly increased in GBM compared to lower grade gliomas (TCGA). **(D)** Immunohistochemistry demonstrates ADAMDEC1 expression in GBM. **(E)** Immunofluorescence shows ADAMDEC1 is absent in tumor-associated microglia (left and center panels; scale bars 50 μm) but is expressed in human xenografted GBM cells (far right panel; scale bar 10 μm). Nuclei are counterstained with DAPI.

We evaluated a role for ADAMDEC1 in maintaining stemness by first enriching GSC and NSCC populations by CD133 expression. Physical separation of the cell types showed increased cellular protein expression of ADAMDEC1 in CD133+ cells compared to their CD133-counterparts across multiple patient-derived cell lines (Fig. 2A). Furthermore, testing conditioned media from GSC and NSCC cultures revealed that GSCs exclusively secrete ADAMDEC1. To evaluate a functional role for ADAMDEC1 in GSC maintenance, we knocked down ADAMDEC1 expression using lentiviral delivery of short hairpin RNA (shRNA) constructs. We found ADAMDEC1 knockdown reduced expression of the stem-cell associated transcription factor SOX2 and increased expression of astrocytic differentiation marker GFAP (Fig. 2B). ADAMDEC1 knockdown also decreased *in vitro* sphere formation (Fig. 2C) and proliferation (Fig. 2D) of primary patient-derived GBM cells compared to nontargeted controls. To further scrutinize the relevance of ADAMDEC1, we orthotopically implanted ADAMDEC1 knockdown cells into immunocompromised mice, and observed a significant increase in survival of tumor-bearing mice compared to controls (Fig. 2E). These data demonstrate ADAMDEC1 is a key regulator of GSCs.

**Figure 2:**
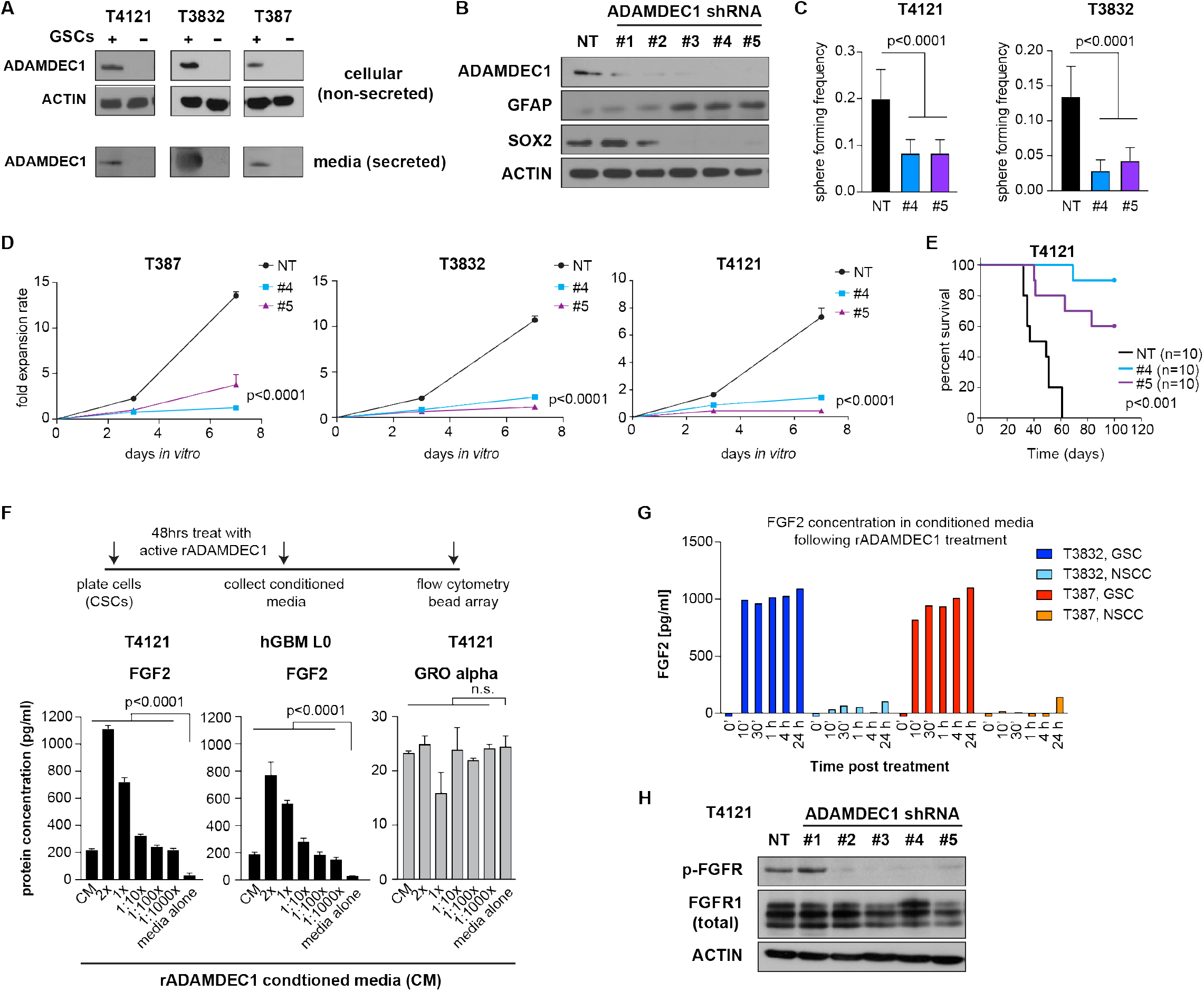
ADAMDEC1 is associated with GBM stemness and secreted by GSCs. **(A)** ADAMDEC1 protein is expressed in GSC, but not in NSCC culture paradigms. Likewise, GSCs secrete ADAMDEC1 into the medium. Depicted are Western blots from cell culture conditioned medium, with 10 μg protein lysate loaded per lane. **(B)** Knockdown of ADAMDEC1 using shRNA. Compared to non-targeting (NT) constructs, ADAMDEC1 knockdown results in decreased SOX2 and increased GFAP expression. **(C)** Sphere-forming frequency is reduced after ADAMDEC1 knockdown (data from two independent experiments, one-way ANOVA). **(D)** ADAMDEC1 knockdown results in decreased cell proliferation in GSC cultures (n=6, non-linear regression). **(E)** Orthotopic implantation of ADAMDEC1 knockdown cells significantly increases survival of tumor-bearing animals compared to control cells (median survival NT=43 d, #4 and #5=100d; n=10 mice/group; log rank test). **(F)** Treatment of GSCs with recombinant ADAMDEC1 results in increased levels of FGF2, but not GRO alpha, in the culture supernatant in a concentration-dependent manner (n=3, two-way ANOVA with Dunnet post-test). **(G)** ELISA shows increased levels of FGF2 in ADAMDEC1-treated GSC, but not in NSCC cultures (data from two independent experiments). **(H)** Western blot showing decreased FGFR phosphorylation after knockdown of ADAMDEC1.

As ADAMDEC1 is a sheddase capable of processing cytokines, we next determined whether ADAMDEC1 promoted GBM growth and progression via cytokine release. We treated GSCs and matching NSCCs with recombinant (r) ADAMDEC1 for 48 hours, after which conditioned media was collected and multiple cytokines were evaluated using anticytokine bead based flow cytometry (Fig. 2F). This experiment showed a dose-dependent increase of soluble FGF2 in the culture media with increasing amounts of rADAMDEC1. In contrast, GRO alpha release was unaffected by rADAMDEC1. We evaluated FGF2 release over time using ELISA to find that GSC, but not NSCC, cultures released FGF2 within minutes following treatment with rADAMDEC1 (Fig. 2G). Finally, ADAMDEC1 knockdown resulted in reduced activation of FGFR signaling, as demonstrated by western blotting using a pan-phospho-FGFR antibody (Fig. 2H).

We next sought to define how FGF2 acts on GSCs by testing whether FGF2 correlated with the GSC-associated transcription factors ZEB1, SOX2, or OLIG2 (26). Using TCGA gene expression data, we found each of these transcription factors correlated strongly with FGF2 (Fig. 3A, S2). To validate these correlations, we used patient-derived GBM cells that had been cultured in EGF only, and treated these cultures with recombinant FGF2. We found that rFGF2 dose-dependently induced expression of ZEB1, SOX2 and OLIG2 (Fig. 3B). In functional assays, rFGF2-treatment increased sphere formation (Fig. 3C). Conversely, a small-molecule inhibitor identified in a screen to block the interaction between FGF2 and FGF receptors (2-Naphthalenesulfonic acid, NSC 65575) (27) reduced sphere formation (Fig. S3) and clonogenicity of patient-derived GBM cells (Fig. 3D). Together, these results implicate FGF2 in GSC activation.

**Figure 3:**
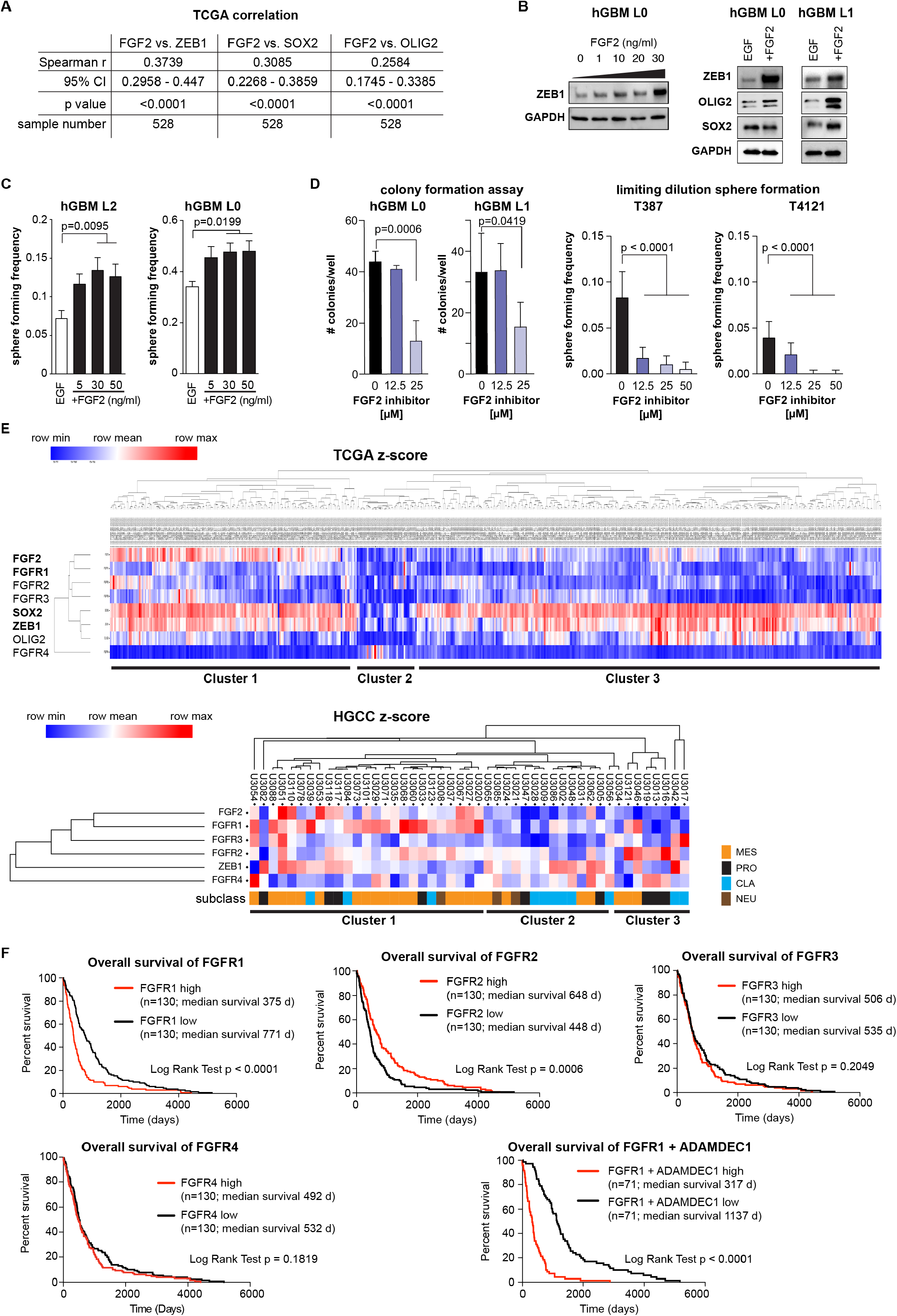
FGF2 promotes stemness in GBM and is linked with FGFR1. **(A)** Spearman correlation of FGF2 with stem cell-associated transcription factors ZEB1, SOX2 and OLIG2 using the Glioblastoma Mutiforme (TCGA, Provisional) Tumor Samples with mRNA data (U133 microarray only) dataset (n=528 samples) shows significant positive correlation for each factor. **(B)** Treatment of primary patient-derived GBM cells with recombinant FGF2 increases ZEB1 expression in a dose-dependent manner. OLIG2 expression is also increased, whereas no change was found for SOX2. **(C)** FGF2 treatment results in increased sphere forming frequency of GSCs in a dose-dependent manner (hGBM L2 n=14, L0 n=10, one-way ANOVA). **(D)** Blocking FGF2 binding to FGFRs using a specific inhibitor (2-Naphthalenesulfonic acid, NSC 65575) reduces colony forming potential of GSCs dose-dependently (n=5, one-way ANOVA). **(E)** Supervised hierarchical clustering of TCGA data (n=528) using FGF2, FGFR1-4, ZEB1, SOX2 and OLIG2 reveals three separate clusters. These clusters could be validated in the HGCC dataset (below) (see text for full description). **(F)** Kaplan-Meier survival analysis of TCGA GBM data, stratified for above-median (high) or below-median (low) gene expression shows a significantly poorer survival in FGFR1 high tumors, but increased survival of FGFR2 high tumors. FGFR3 expression has no effect on survival. Combined analysis of FGFR1 and ADAMDEC1 shows a very strong effect on survival.

FGF2 can bind to FGFR1-4, therefore we used a bioinformatic approach to determine whether any FGFRs show association with GBM patient outcome. We performed hierarchical cluster analysis of TCGA data using FGF2, FGFR1-4, ZEB1, SOX2 and OLIG2 as determinants. This revealed the existence of three principal clusters (Fig. 3E), with cluster 1 comprising of ~30% of samples and showing association of FGF2, FGFR1-3 and stem cell transcription factors (ZEB1, OLIG2, and SOX2). Cluster 2 was the smallest and showed low expression of FGF2, FGFR1-3 and stem cell transcription factors. Conversely, this was the only cluster where FGFR4 was present. Cluster 3 constitutes most (>50%) of the samples, and shows high expression of ZEB1, SOX2, and OLIG2, but not of FGF2 or its receptors. Gene set enrichment analysis revealed an overlap of cluster 1 with signatures of the classical and mesenchymal molecular subclasses (4), while cluster 2 showed positive correlation with mesenchymal and negative correlation with all other signatures, and cluster 3 was enriched for neural and proneural subclass signatures (Fig. S4).

Because data in the TCGA repository may be affected by the presence of non-tumor cells in patient specimens, we validated our cluster analysis in the HGCC dataset, which is derived from patient cell lines (28). Consistently, we found all three clusters represented. In this dataset, FGFR1 was strongest associated with cluster 1, which was also most enriched for the mesenchymal subclass.

This demonstrates an association of FGF2 and stemness-associated transcription factors in a significant fraction of GBM specimens and that considerable heterogeneity exists among these samples.

We used Kaplan-Meier analysis of the TCGA Glioblastoma Multiforme (provisional) dataset to further investigate which FGFR may be most relevant for patient outcome. This analysis confirmed that high expression of FGFR1 is associated with poor outcome, while for FGFR2 the relationship was inverse of this with augmented FGFR2 expression correlating to a better prognosis (Fig. 3F). No significant correlation was apparent for FGFR3 and patient survival. Of note, combinatorial analysis for FGFR1 and ADAMDEC1 showed a strong association with survival.

We next determined the FGFR expression in patient-derived GBM cells and found that FGFR1-3 were expressed, but that FGFR4 was consistently absent (Fig. 4A), as suggested by our cluster analyses (Fig. 3E). Using flow cytometry to quantify FGFR1-3 expression on GBM cells (Fig. 4B, S5A), we found FGFR1 present on a small subset of cells (1-5%), consistent with a stem cell population. By contrast, FGFR2 was expressed on a large fraction (30-50%) of cells. FGFR3 expression showed large variance, ranging from 5-25%. To test the functional relationship of different FGFR subtypes with GSCs, we performed lentiviral shRNA-mediated knockdown of FGFR1-3 in patient-derived GBM cells (Fig. 4C, S5B). We found that only FGFR1 loss resulted in a decrease of sphere forming capacity (Fig. 4D), which was accompanied by a decrease in ZEB1, SOX2, and OLIG2 expression (Fig 4E), and a decrease in proliferation (Fig. S5C). Loss of FGFR1, but not of FGFR2 or FGFR3, abolished the FGF2-mediated increase in self-renewal (Fig. S6). Importantly, FGFR1 knockdown increased survival of tumor-bearing mice after orthotopic transplantation (Fig. 4F). To further substantiate this pivotal role of FGFR1 in GSCs, we performed rescue experiments. Targeted expression of full-length FGFR1 increased expression of ZEB1, SOX2 and OLIG2 in control cells, and restored ZEB1, SOX2 and OLIG2 expression in FGFR1 knockdown cells (Fig. 4G). Concomitantly, over-expression of FGFR1 increased sphere formation in control and FGFR1 knockdown cells (Fig. 4H). By contrast, ZEB1 knockdown negated the effects of FGFR1 over-expression on expression of SOX2 and OLIG2, as well as on sphere formation, indicating that ZEB1 is downstream of FGFR1.

**Figure 4:**
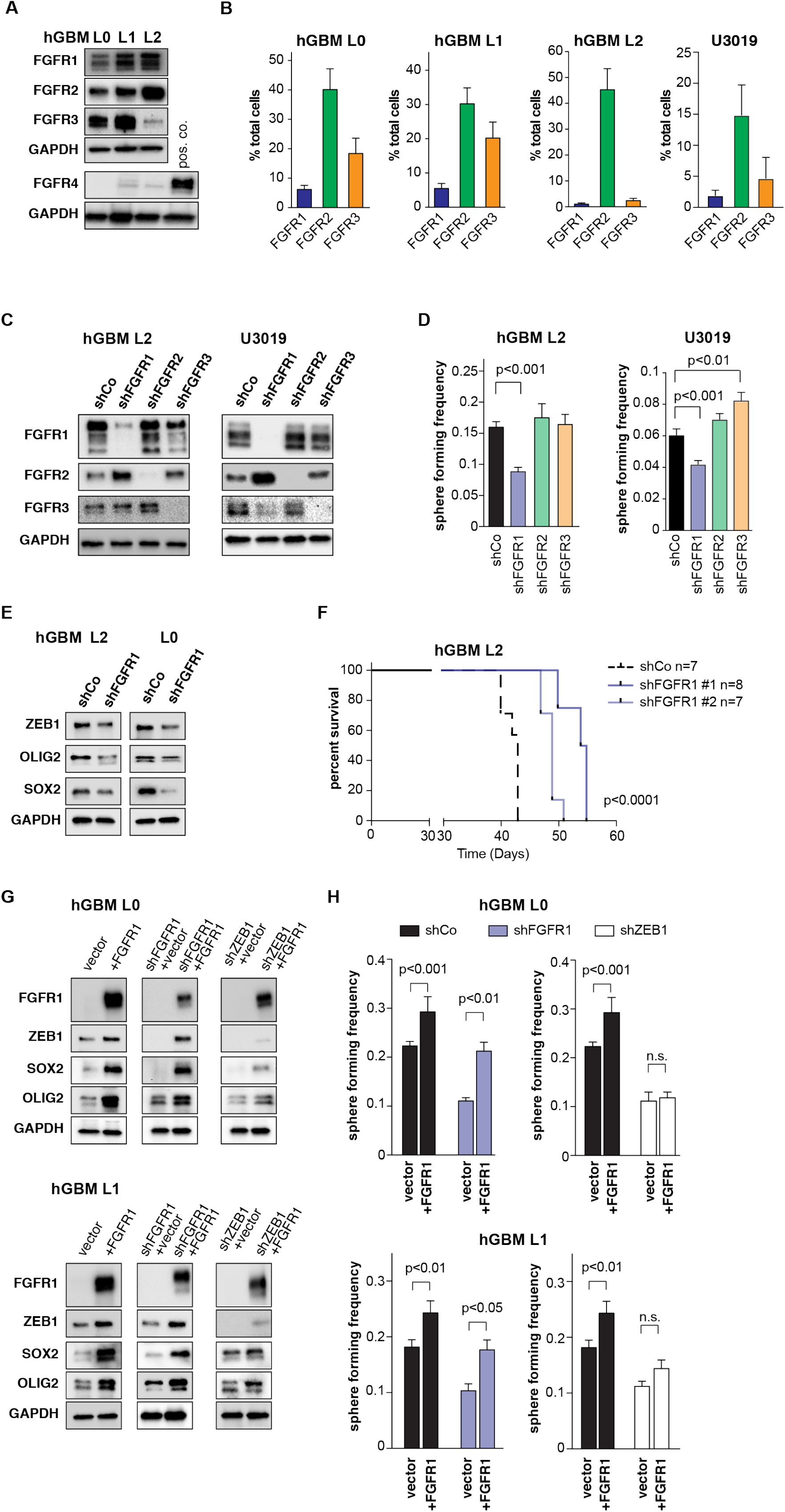
FGF2 promotes stemness in GBM through FGFR1. **(A)** Western blotting shows expression of FGFR1-3, but not FGFR-4 in primary patient-derived GBM cell lines (hGBM L0, L1, L2). **(B)** Flow cytometry quantification of FGFR1-3 expression in patient-derived human GBM lines (n=3 independent experiments/sample). Note that FGFR1 is expressed in a small subset of each line. **(C)** Knockdown of FGFR1-3 using shRNA constructs shows specificity for each receptor. Additional constructs are shown in Fig. S5. **(D)** Knockdown of FGFR1, but not FGFR2 or FGFR3, results in decreased sphere-forming frequency compared to control cells (hGBM L2 n=6, U3019 n=9, one-way ANOVA). **(E)** FGFR1 knockdown decreases expression of ZEB1, SOX2 and OLIG2 in GSCs. **(F)** Orthotopic implantation of FGFR1 knockdown cells significantly increases survival of tumor-bearing animals (median survival shCo: 43d, shFGFR1#1: 54.5d, shFGFR1#2: 49d; log-rank test). **(G)** Expression of full-length FGFR1 increases ZEB1 expression in control cells, and rescues ZEB1 expression in FGFR1 knockdown cells. **(H)** Full-length FGFR1 expression increases sphere-forming frequency in control cells (black bars), and rescues sphere-forming frequency of FGFR1 knockdown cells to control levels (blue bars). Knockdown of ZEB1 negates the effect of FGFR1 expression (white bars) (n=9, two-way ANOVA).

As FGFR1 is functionally relevant for stem cell maintenance in GBM, we hypothesized that this receptor may identify a GSC population. To test this hypothesis, we first quantified FGFR1 expression under culture conditions conducive to GSC maintenance (sphere cultures supplemented with EGF) or differentiation (adherent cultures with growth factor withdrawal and 10% serum). We found that FGFR1 expression is high in GSC cultures, whereas FGFR1 levels decrease under differentiation conditions (Fig. 5A). The expression of the stem cell transcription factors ZEB1, SOX2 and OLIG2 followed the same pattern. By contrast, FGFR2 expression increased in differentiation conditions (Fig. 5A). To test whether FGFR1 may be a marker of GSCs, we isolated FGFR1-expressing cells via flow cytometry (Fig. 5B). Using an independent FGFR1-antibody, we confirmed increased FGFR1 expression in the FGFR1+ fraction. Likewise, expression of ZEB1, SOX2 and OLIG2 were enriched in the FGFR1+ fraction (Fig. 5B). Next, we tested FGFR1+ cells in functional assays for stemness immediately after FACS. We found that FGFR1+ cells showed greater colony forming potential than FGFR1-cells (Fig. 5C). Finally, we used limiting dilution orthotopic xenografts to determine the stem cell frequency of FGFR1 + and FGFR1-cells (Fig. 5D). Extreme limiting dilution analysis (29) of tumor formation in this paradigm demonstrated an approximate 5-fold enrichment of GSCs in FGFR1+ cells. Together, these data strongly support that (i) FGFR1 transduces FGF2 signal to induce GSC self-renewal, (ii) ZEB1, SOX2 and OLIG2 are downstream targets of FGF2/FGFR1 signaling, and (iii) FGFR1 is a surface marker of GSCs.

**Figure 5:**
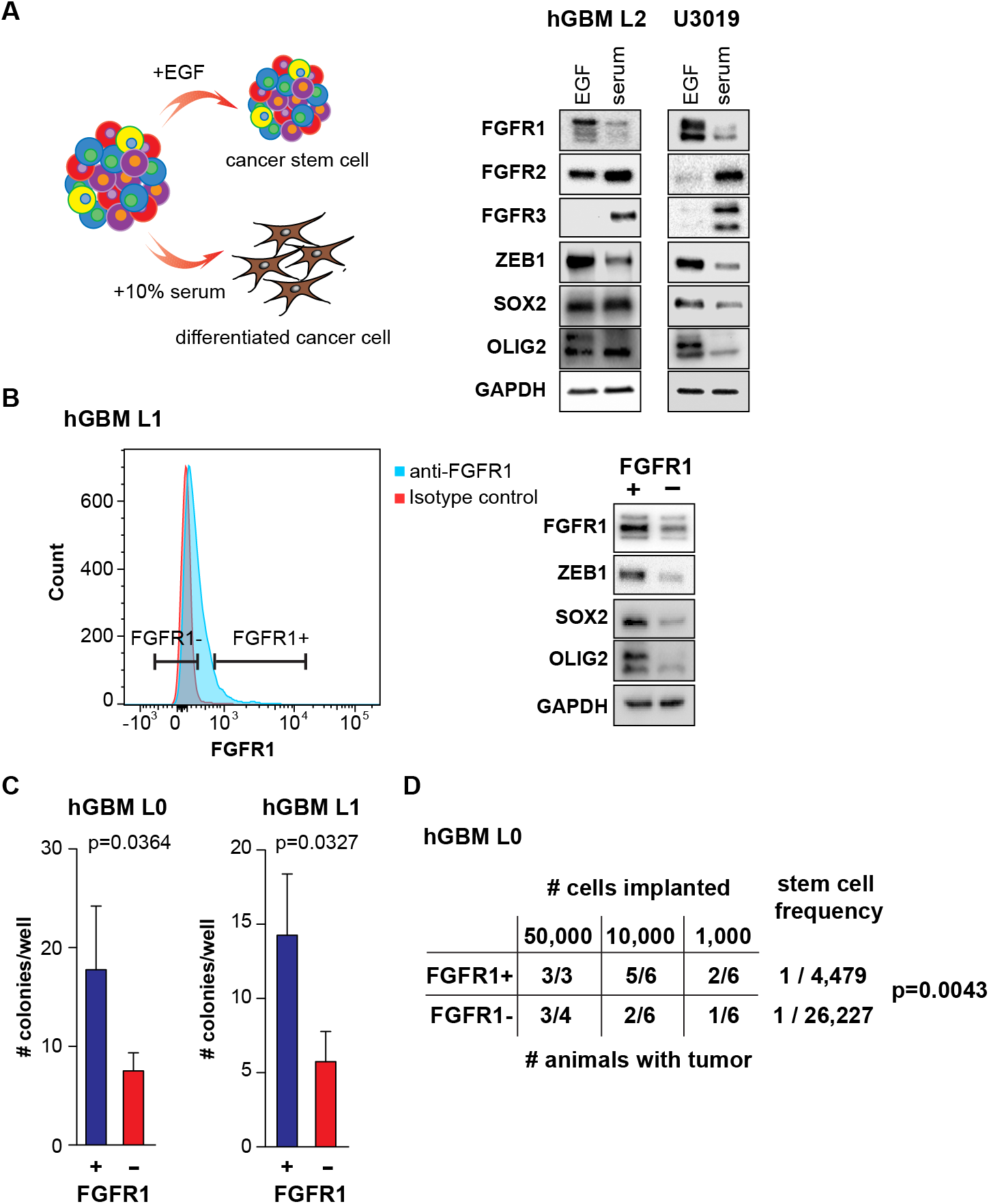
FGFR1 is endogenously associated with a stem cell population. **(A)** Expression of FGFR1, FGFR2, ZEB1, and SOX2 is affected by the culture paradigm. FGFR1, ZEB1 and SOX2 expression are higher in GSC conditions, whereas FGFR2 and FGFR3 increase upon differentiation. **(B)** Flow cytometry isolation of FGFR1 + cells. Histogram shows positive FGFR1 staining in GBM cells compared to isotype control. Additional plots are shown in Fig. S5. Western analysis using an independent FGFR1 antibody demonstrates higher FGFR1 expression in FGFR1 + cells post sort, as well as increased ZEB1, SOX2 and OLIG2 expression. **(C)** FGFR1+ cells show greater potential for colony formation in a collagen matrix. Cells were plated in colony forming assays immediately after sorting (paired t-test). **(D)** Limiting dilution orthotopic xenografts reveal greater tumorigenicity of FGFR1+ cells. Stem cell frequency was calculated using ELDA (Chi square test).

Our results indicate that GSCs secrete ADAMDEC1, causing release of FGF2 that induces GSC maintenance via FGFR1 signaling to ZEB1. We next sought to address how the expression of ADAMDEC1 in GSCs might be regulated. Speculating that ZEB1 may induce ADAMDEC1 expression, we tested whether ADAMDEC1 forms a positive feedback loop in GSCs with FGFR1 and ZEB1. Indeed, FGF2 treatment induced expression of ADAMDEC1 compared to EGF stimulation (Fig. 6A). We tested whether this effect on ADAMDEC1 expression was mediated by FGFR1, and found that FGFR1 knockdown reduced ADAMDEC1 expression, while targeted expression of FGFR1 increased ADAMDEC1 levels (Fig. 6B). Finally, we demonstrated that ZEB1 knockdown also decreases ADAMDEC1 protein expression (Fig. 6C).

**Figure 6:**
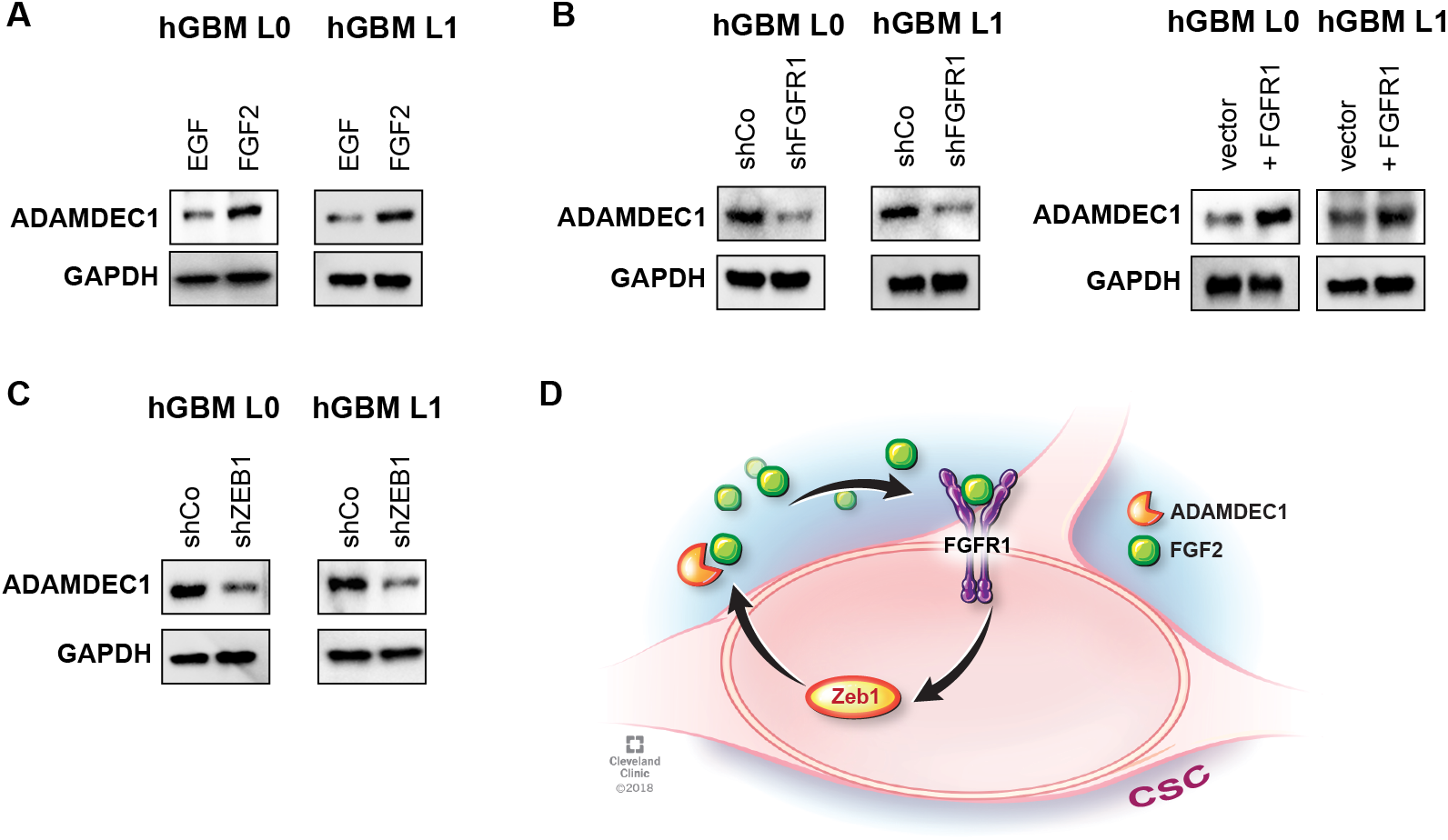
FGFR1 regulates ADAMDEC1 through ZEB1. **(A)** Western analysis shows increased ADAMDEC1 expression after GSC treatment with FGF2. **(B)** ADAMDEC1 expression is decreased after FGFR1 knockdown and increased after FGFR1 expression in GSCs. **(C)** ZEB1 knockdown results in decreased expression of ADAMDEC1. **(D)** Diagram depicting the ADAMDEC1-FGFR1-ZEB1 feedback loop.

Together, our data supports the existence of a positive feedback loop that activates stemness in GBM (Fig. 6D). ADAMDEC1 is secreted by GSCs, and releases FGF2 from the extracellular matrix in the tumor microenvironment. FGF2 binds to FGFR1 on the surface of GSCs, where activation of this receptor leads to increased expression of the downstream targets ZEB1, SOX2 and OLIG2. ZEB1 mediates FGF2/FGFR1 effects on stemness, and regulates expression of ADAMDEC1, thereby completing the loop.

## Discussion

The mechanisms by which GSCs maintain their stemness across different niches in the tumor landscape, including hypoxic, vascular, invasive niches, is incompletely understood and a matter of intense investigation (14). Our study identifies a new feedback loop that enables GSCs to access FGF2 in the tumor microenvironment through secretion of the ADAMDEC1. ADAMDEC1 is a novel member of the ADAM family of metalloproteinases due to the absence of a transmembrane domain and altered catalytic domain, features shared by no other ADAM family member (20, 30, 31), which result in a unique secreted, soluble protease with distinct ligand specificity (20, 21). Additionally, ADAMDEC1 was shown to modulate apical membrane extrusion of epithelial cells (32). Here, we demonstrate ADAMDEC1 is highly expressed in glioma with enhanced expression correlating to increasing with tumor grade, and ADAMDEC1 expression correlated with patient survival. Further, we demonstrate that ADAMDEC1 was enriched in GSC populations and regulated cell proliferation, sphere formation and tumorigenesis. ADAMDEC1 was recently shown to mediate the cleavage and release of active EGF, an important GSC trophic factor (21), although in our data FGF2 had a stronger effect on GSC stemness than EGF alone.

FGF2 binds to all four members of the FGF receptor family, with splice isoforms mediating binding affinity (33). FGF2/FGFR1 signaling promotes glioma growth and radioresistance (34, 35) and higher FGFR1 expression is associated with poor outcome for these tumors (36). A recent study indicated a link between FGFR1 and ZEB1 in GBM (37), but definitive evidence for FGFR1 regulating stemness including limiting-dilution *in vivo* transplantation was still lacking, which are presented in our study. By contrast, FGFR2 expression is reduced in GBM compared to low-grade glioma (38), and higher FGFR2 levels are associated with improved survival (39). FGFR3 was the second-most differentially expressed gene between infiltrating and tumor-core GBM cells in a recent single-cell study (40), and FGFR3-TACC3 gene fusions have been identified as oncogenic drivers of GBM growth (41).

FGF2 activates FGFR1 on the GSC cell surface, which in turn induced expression of stem cell transcription factors. Crucially, FGFR1 was the only FGFR that we found to be functionally associated with tumor cell sphere formation in culture, a hallmark of stemness. The relevance of FGFR1 for GSCs is further supported by increased survival after loss of FGFR1, as well as increased tumorigenicity of FGFR1+ cells. We and others have shown that ZEB1 is a key regulator of stemness, invasion and chemoresistance in GBM (7, 12, 42), and consistently we find that ZEB1 mediates the stemness effects of FGF2/FGFR1 signaling. Singh and colleagues have recently demonstrated that ZEB1, SOX2 and OLIG2 form an autonomous transcriptional loop in GBM, and can regulate their expression reciprocally (26). Our data supports this, as the FGF2/FGFR1 complex activates expression of SOX2 and OLIG2 in our system as well, and ZEB1, SOX2, and OLIG2 are linked in hierarchical cluster analysis of TCGA data. It is tempting to speculate due to the apparent co-dependency of the transcription factors ZEB1, SOX2 and OLIG2 that interference with any one member of this circuit may disrupt stemness in GSCs. Our data indicate this may be the case, as loss of ZEB1 is sufficient to abrogate the effects of FGF2 stimulation and increased FGFR1 expression on GSCs.

We additionally found that ZEB1 regulates expression of ADAMDEC1, creating a feedback loop that would enable GSCs to thrive in the CNS, as FGF2 is highly prevalent across the brain and in the cerebrospinal fluid. We find that in the TCGA dataset, approximately 30% of GBM patients are characterized by increased expression of FGF2, FGFRs, and ZEB1, and thus may benefit from therapeutic intervention aimed at disrupting this feedback loop. Importantly, we show that pharmacological intervention of FGF2 binding to its cognate receptors can block stemness in GSCs, indicating that this may be an exploitable fulcrum for future therapies. This may be particularly relevant for those patients, where cluster analysis shows higher expression of components of this feedback loop. Currently, clinical trials are underway for FGF receptor tyrosine kinase inhibitors (43). While these compounds are selective for FGFR1-3 over FGFR4, no compounds exist at present that are selective for FGFR1 over FGFR2-4. Furthermore, trials for FGFR inhibitors in glioma only recruit patients with amplification or mutation of FGFR genes, which constitute only approximately 3-5% of all GBM patients. Our data indicate that a much larger fraction of patients could benefit from anti-FGFR therapies, should such trials be successful.

Taken as a whole, our data identify a novel GSC signaling axis: ADAMDEC1-FGF2-FGFR1-ZEB1 and present a druggable, translational point of fragility.

## Methods

### Primary human glioblastoma cells

Human glioblastoma (hGBM) cells were cultured as described previously (7, 12). Briefly, for GSC sphere culture, 5×10^4^ cells/ml were plated in N2 medium (Thermo Fisher) supplemented with 2% bovine serum albumin (Fisher Scientific) containing 20 ng/ml recombinant human EGF (Peprotech). For some experiments, recombinant human FGF2 (Peprotech) was added at concentrations ranging from 5 ng/ml to 80 ng/ml. For differentiation experiments, spheres were dissociated into single cells and plated in DMEM/F12 (Thermo Fisher) supplemented with 10% fetal bovine serum (FBS).

Cell lines from the HGCC repository (28) were cultured in Neurobasal medium (Thermo Fisher) supplemented with B27 (Thermo Fisher) and 20ng/ml EGF and FGF2. Spheres were passaged when they reached an average diameter of 150 μm.

In some experiments, previously established GBM xenografts obtained from Duke University and the University of Florida were used and maintained as previously described (44, 45). Tissue was digested with papain (Worthington) as previously described (45) and dissociated cells allowed to recover overnight prior to use. Thereafter, dissociated cells were sorted based on CD133 expression using magnetic beads (Miltenyi). CD133-positive CSCs were maintained in Neurobasal medium (Life Technologies) supplemented with penicillin/streptomycin (50 U/ml final concentration), L-glutamine (2 mM), B27 (Life Technologies), sodium pyruvate (1 mM), EGF (20 ng/ml, R&D Systems) and FGF2 (20 ng/ml, R&D Systems). CD133-negative NSCCs were cultured in DMEM supplemented with 10% FBS and penicillin/streptomycin (50 U/ml).

### Plasmids and lentiviral transduction

Control and knockdown ZEB1, FGFR1, FGFR2 and FGFR3 plasmids were purchased from Dharmacon. Control and knockdown ADAMDEC1 plasmids were purchased from Sigma. Different clones for each shRNA plasmid were tested and the best knockdowns were selected to produce lentiviral particles. A plasmid for over expression of FGFR1 was a gift from Dominic Esposito (Addgene plasmid #70367) and cloned into an expression vector (pHIV-IRES-mRFP) using the Gateway system (Invitrogen). Lentiviral particles were generated by co-transfecting HEK293T cells with second generation packaging plasmids (psPAX2 and pMD2.G) using Lipofectamine 3000 (Invitrogen). Medium containing lentiviral particles was collected 48 h and 72 h after transfection. Viral supernatants were combined and filtered with a 0.45 μm pore size filter, followed by ultracentrifugation at 185,000 rcf with a L8-70M Ultracentrifuge (Beckman). Pelleted viral particles were diluted in 200 μl of N2 medium. Concentrated viral particles were aliquoted and stored at −80°C.

For lentiviral transduction, 1×10^5^ cells were pre-incubated for 1-2 h in N2 medium without antibiotics and 1 μg/μl of polybrene (Santa Cruz) to increase transduction efficiency. 18h after transduction, medium was replaced with complete N2 medium and growth factors.

### FGF2 inhibitors

The small molecule inhibitors of FGF2, NSC-47762, NSC-58057, NSC-65575 and NSC-65576 (27) were obtained from the Developmental Therapeutics Program (DTP), division of Cancer Treatment and Diagnosis, NCI, NIH (USA).

### Sphere-forming frequency assays

Limiting dilution analysis was carried out, following 14 days’ incubation, in a 96-well format with 24 wells of each dilution; 1, 5, 10, and 20 cells/well. Cells were sorted using the BD FACS ARIA II Flow Cytometer. Limiting dilution plots and stem cell frequencies were calculated using ELDA analysis (http://bioinf.wehi.edu.au/software/elda/index.html) (29). Sphere-forming assays were performed as described (7), with the following modifications: 100-200 single cells were seeded per well in 96 well plates, with 6 replicates per condition. Cells were cultured in 80μl of N2 medium supplemented with 20ng/ml of EGF and/or 30ng/ml of FGF2 as specified in the text. The number of spheres with a diameter greater than 70 μm was quantified on a GelCount analyzer (Oxford Optronix) 5 days after seeding.

### Clonogenicity assays

For colony-forming cell assays, 1×10^4^ hGBM single cells were cultured in N2 supplemented with 20ng/ml of EGF, 30ng/ml FGF2 and collagen (Stem Cell Technologies) at a ratio of 1:3 (collagen/medium). Cells were cultured at 37°C and supplemented with fresh growth factors twice/week. Colonies greater than 200μm of diameter were counted two weeks after plating on a GelCount analyzer.

### Flow cytometry and cell sorting

Immunostaining and flow cytometry was performed as described (12). Data was acquired on a BD LSR Fortessa (BD Bioscience), using FACSDIVA software (BD Bioscience) and analyzed with a FlowJo ver. 8.8.7 (Tree Star, Inc). For cell sorting, stained single cell suspensions were purified on a BD FACSAria Fusion (BD Bioscience) and immediately used for downstream experiments.

### Protein isolation and Western blotting

Protein isolation and quantification was performed as described (7, 21). For Western blotting, 5 to 20 μg of sample were mixed with equal volume of 2x Laemmli buffer (Bio-Rad) containing β-mercaptoethanol, and denatured at 95°C for 5 min. Following separation on a mini-protean 4-15% Bis-Tris gel (Bio-Rad), proteins were transferred onto PVDF membranes using a Mini Trans-Blot Turbo Transfer System (Bio-Rad). Membranes were blocked and probed for primary and secondary antibodies as described (7) (see Table S1 for antibody information), and visualized using Clarity Western ECL substrate (Bio-Rad) on a ChemiDoc MP imaging system (Biorad). Results were normalized to GAPDH or ACTNB as housekeeping genes.

### Animal experiments

Female SCID mice aged 4-6 weeks were used for orthotopic xenografts. Animal care and handling, and all procedures were performed according to NIH, FELASA and institutional guidelines and approved by the UK home office (PPL30/3331) and the Institutional Animal Care and Use Committee of the Cleveland Clinic Foundation (protocol 2012-0752). Intracranial tumor transplants were performed as described previously (7, 46). Depending on the experiment, 1,000 – 50,000 cells were stereotactically implanted in 5 μl of media devoid of growth factors/supplements. Mice were maintained under Isoflurane anesthesia during procedures. Mice were monitored daily for the development of neurological signs and body weight loss. Animals at endpoint were transcardially perfused using 2% paraformaldehyde and the brains removed for histology.

### Tissue preparation and immunofluorescence

Post-fixed brains were cryoprotected, embedded and 30 μm coronal sections prepared as described (47). Sections were prepared for immunofluorescence staining, mounted and coverslipped using standard protocols.

### Bioinformatics

TCGA, NCI, Gravendeel, Murat and Kamoun mRNA datasets were analyzed via the online tool GlioVis (http://gliovis.bioinfo.cnio.es/). Further analysis of survival correlations was performed using the Xena platform (https://xena.ucsc.edu/welcome-to-ucsc-xena/).

### Hierarchical cluster analysis

HGCC cell line expression data (28) was clustered using the hierarchical clustering module (48) from GenePattern (https://cloud.genepattern.org) (49) with Pearson correlation as distance measure, using row centering and normalization. Z-Score values from the Glioblastoma Mutiforme (TCGA, Provisional) Tumor Samples with mRNA data (U133 microarray only) (528 samples) data set was downloaded from cBioportal (50) and clustered with Pearson correlation as distance measure, using the same GenePattern module as before.

Gene Set Enrichment Analysis was performed on our identified clusters using the GSEA module from GenePattern (51), this was done using the Glioblastoma Mutiforme (TCGA, Provisional) Tumor Samples with mRNA data (U133 microarray only) data set downloaded from UCSC Xena Platform website (https://www.biorxiv.org/content/early/2018/05/18/326470). Clusters were compared one versus the rest using the Verhaak Glioblastoma Proneural, Classical, Neural and Mesenchymal (4) gene sets from MsigDB (52). The results for the TCGA data set here are in whole or part based upon data generated by the TCGA Research Network: http://cancergenome.nih.gov/.

### Image acquisition

Images were acquired using a Leica TCS-SP8-AOBS inverted confocal microscope (Leica Microsystems).

### Statistical analysis

Statistical analyses were performed in GraphPad Prism version 7, using statistical tests as indicated in the text. In all analyses, p values <0.05 were deemed as significant.

## Acknowledgements

Funding was provided by Medical Research Council grant (MR/S007709/1), Tenovus Cancer Care (TIG2015/L19) and a Cancer Research UK Cardiff Centre Development Fund Award to FAS and the Lisa Dean Moseley Foundation to JDL and TMM.

## Supplementary Information

**Fig. S1: (A)** Kaplan-Meier analysis of GBM patient survival after stratification for ADAM family members in the LeeY dataset. **(B)** Kaplan-Meier analysis of patient survival (combined high-grade and low-grade glioma) after stratification for ADAMDEC1 in the TCGA Glioma dataset.

**Fig. S2:** Spearman correlations of FGF2 and ZEB1 (left), SOX2 (middle) and OLIG2 (right) in the TCGA dataset, based on expression z-scores.

**Fig. S3:** Sphere-forming assay using three compounds that have been shown to block binding FGF2 to its receptors. Bar graphs show sphere-forming frequency relative to vehicle control (DMSO). NSC 65575 was the only compound effective in this assay and was tested in subsequent colony-forming assays.

**Fig. S4:** Gene-set enrichment analysis for clusters 1-3 from hierarchical cluster analysis of TCGA data in Fig. 3, using molecular subclasses as defined by Verhaak et al. 2010.

**Fig. S5: (A)** FGFR1 antibody specificity in flow cytometry. Shown are histograms for staining intensity (x-axis) versus event counts (y-axis) for isotype control, FGFR1-stained samples, and FGFR1 knockdown samples. Note the lack of staining after FGFR1 knockdown. **(B)** Gating strategy for FACS sorting of FGFR1 + and FGFR1-cells. **(C)** Western blot demonstrates specificity of FGFR knockdown constructs. Shown are multiple constructs for FGFR1, −2 and −3. In all cases construct #1 was selected for downstream experiments. **(D)** Loss of FGFR1, but not FGFR2 or −3, affects proliferation in EGF (hGBM L1) and EGF/FGF2 (U3019) treated cells. Bars represent fold-change of expansion rates (n=3 per cell line; p<0.05, one-way ANOVA).

**Fig. S6: (A)** Sphere-forming assays with EGF or EGF+FGF2 reveal that knockdown of FGFR1 consistently blocks self-renewal in two patient-derived GBM cell lines. By contrast, loss of FGFR2 increased self-renewal (hGBM L2), or showed no effect (hGBM L1) in response to FGF2 stimulation. Loss of FGFR3 showed inconsistent effects. **(B)** Loss of ZEB1 also blocked FGF2-induced self-renewal, supporting that ZEB1 is downstream of FGFR1, and sufficient to mediate self-renewal effects of FGF2.

**Table S1:** List of antibodies used in this study.

## References

1. Stupp R, Hegi ME, Mason WP, van den Bent MJ, Taphoorn MJ, Janzer RC, et al. Effects of radiotherapy with concomitant and adjuvant temozolomide versus radiotherapy alone on survival in glioblastoma in a randomised phase III study: 5-year analysis of the EORTC-NCIC trial. Lancet Oncol. 2009;10:459–66.

2. Weller M, Butowski N, Tran DD, Recht LD, Lim M, Hirte H, et al. Rindopepimut with temozolomide for patients with newly diagnosed, EGFRvIII-expressing glioblastoma (ACT IV): a randomised, double-blind, international phase 3 trial. Lancet Oncol. 2017;18:1373–85.

3. Wick W, Gorlia T, Bendszus M, Taphoorn M, Sahm F, Harting I, et al. Lomustine and Bevacizumab in Progressive Glioblastoma. N Engl J Med. 2017;377:1954–63.

4. Verhaak RG, Hoadley KA, Purdom E, Wang V, Qi Y, Wilkerson MD, et al. Integrated genomic analysis identifies clinically relevant subtypes of glioblastoma characterized by abnormalities in PDGFRA, IDH1, EGFR, and NF1. Cancer Cell. 2010;17:98–110.

5. Capper D, Jones DTW, Sill M, Hovestadt V, Schrimpf D, Sturm D, et al. DNA methylation-based classification of central nervous system tumours. Nature. 2018;555:469–74.

6. Chen J, Li Y, Yu TS, McKay RM, Burns DK, Kernie SG, et al. A restricted cell population propagates glioblastoma growth after chemotherapy. Nature. 2012;488:522–6.

7. Siebzehnrubl FA, Silver DJ, Tugertimur B, Deleyrolle LP, Siebzehnrubl D, Sarkisian MR, et al. The ZEB1 pathway links glioblastoma initiation, invasion and chemoresistance. EMBO molecular medicine. 2013;5:1196–212.

8. Bao S, Wu Q, McLendon RE, Hao Y, Shi Q, Hjelmeland AB, et al. Glioma stem cells promote radioresistance by preferential activation of the DNA damage response. Nature. 2006;444:756–60.

9. Baysan M, Woolard K, Cam MC, Zhang W, Song H, Kotliarova S, et al. Detailed longitudinal sampling of glioma stem cells in situ reveals Chr7 gain and Chr10 loss as repeated events in primary tumor formation and recurrence. International journal of cancer. 2017;141:2002–13.

10. Singh SK, Clarke ID, Terasaki M, Bonn VE, Hawkins C, Squire J, et al. Identification of a cancer stem cell in human brain tumors. Cancer research. 2003;63:5821–8.

11. Vanner RJ, Remke M, Gallo M, Selvadurai HJ, Coutinho F, Lee L, et al. Quiescent sox2(+) cells drive hierarchical growth and relapse in sonic hedgehog subgroup medulloblastoma. Cancer Cell. 2014;26:33–47.

12. Hoang-Minh LB, Siebzehnrubl FA, Yang C, Suzuki-Hatano S, Dajac K, Loche T, et al. Infiltrative and drug-resistant slow-cycling cells support metabolic heterogeneity in glioblastoma. EMBO J. 2018.

13. Lan X, Jorg DJ, Cavalli FMG, Richards LM, Nguyen LV, Vanner RJ, et al. Fate mapping of human glioblastoma reveals an invariant stem cell hierarchy. Nature. 2017;549:227–32.

14. Lathia JD, Heddleston JM, Venere M, Rich JN. Deadly teamwork: neural cancer stem cells and the tumor microenvironment. Cell Stem Cell. 2011;8:482–5.

15. Dong F, Eibach M, Bartsch JW, Dolga AM, Schlomann U, Conrad C, et al. The metalloprotease-disintegrin ADAM8 contributes to temozolomide chemoresistance and enhanced invasiveness of human glioblastoma cells. Neuro Oncol. 2015;17:1474–85.

16. Musumeci G, Magro G, Cardile V, Coco M, Marzagalli R, Castrogiovanni P, et al. Characterization of matrix metalloproteinase-2 and -9, ADAM-10 and N-cadherin expression in human glioblastoma multiforme. Cell Tissue Res. 2015;362:45–60.

17. Siney EJ, Holden A, Casselden E, Bulstrode H, Thomas GJ, Willaime-Morawek S. Metalloproteinases ADAM10 and ADAM17 Mediate Migration and Differentiation in Glioblastoma Sphere-Forming Cells. Mol Neurobiol. 2017;54:3893–905.

18. Wolpert F, Tritschler I, Steinle A, Weller M, Eisele G. A disintegrin and metalloproteinases 10 and 17 modulate the immunogenicity of glioblastoma-initiating cells. Neuro Oncol. 2014;16:382–91.

19. Sarkar S, Zemp FJ, Senger D, Robbins SM, Yong VW. ADAM-9 is a novel mediator of tenascin-C-stimulated invasiveness of brain tumor-initiating cells. Neuro Oncol. 2015;17:1095–105.

20. Lund J, Troeberg L, Kjeldal H, Olsen OH, Nagase H, Sorensen ES, et al. Evidence for restricted reactivity of ADAMDEC1 with protein substrates and endogenous inhibitors. J Biol Chem. 2015;290:6620–9.

21. Chen R, Jin G, McIntyre TM. The soluble protease ADAMDEC1 released from activated platelets hydrolyzes platelet membrane pro-epidermal growth factor (EGF) to active high-molecular-weight EGF. J Biol Chem. 2017;292:10112–22.

22. Haley EM, Kim Y. The role of basic fibroblast growth factor in glioblastoma multiforme and glioblastoma stem cells and in their in vitro culture. Cancer letters. 2014;346:1–5.

23. Bian XW, Du LL, Shi JQ, Cheng YS, Liu FX. Correlation of bFGF, FGFR-1 and VEGF expression with vascularity and malignancy of human astrocytomas. Anal Quant Cytol Histol. 2000;22:267–74.

24. Pollard SM, Yoshikawa K, Clarke ID, Danovi D, Stricker S, Russell R, et al. Glioma stem cell lines expanded in adherent culture have tumor-specific phenotypes and are suitable for chemical and genetic screens. Cell Stem Cell. 2009;4:568–80.

25. Bowman RL, Wang Q, Carro A, Verhaak RG, Squatrito M. GlioVis data portal for visualization and analysis of brain tumor expression datasets. Neuro Oncol. 2017;19:139–41.

26. Singh DK, Kollipara RK, Vemireddy V, Yang XL, Sun Y, Regmi N, et al. Oncogenes Activate an Autonomous Transcriptional Regulatory Circuit That Drives Glioblastoma. Cell Rep. 2017;18:961–76.

27. Foglieni C, Pagano K, Lessi M, Bugatti A, Moroni E, Pinessi D, et al. Integrating computational and chemical biology tools in the discovery of antiangiogenic small molecule ligands of FGF2 derived from endogenous inhibitors. Sci Rep. 2016;6:23432.

28. Xie Y, Bergstrom T, Jiang Y, Johansson P, Marinescu VD, Lindberg N, et al. The Human Glioblastoma Cell Culture Resource: Validated Cell Models Representing All Molecular Subtypes. EBioMedicine. 2015;2:1351–63.

29. Hu Y, Smyth GK. ELDA: extreme limiting dilution analysis for comparing depleted and enriched populations in stem cell and other assays. J Immunol Methods. 2009;347:70–8.

30. Lund J, Olsen OH, Sorensen ES, Stennicke HR, Petersen HH, Overgaard MT. ADAMDEC1 is a metzincin metalloprotease with dampened proteolytic activity. J Biol Chem. 2013;288:21367–75.

31. Bates EE, Fridman WH, Mueller CG. The ADAMDEC1 (decysin) gene structure: evolution by duplication in a metalloprotease gene cluster on chromosome 8p12. Immunogenetics. 2002;54:96–105.

32. Yako Y, Hayashi T, Takeuchi Y, Ishibashi K, Kasai N, Sato N, et al. ADAM-like Decysin-1 (ADAMDEC1) is a positive regulator of Epithelial Defense Against Cancer (EDAC) that promotes apical extrusion of RasV12-transformed cells. Sci Rep. 2018;8:9639.

33. Zhang X, Ibrahimi OA, Olsen SK, Umemori H, Mohammadi M, Ornitz DM. Receptor specificity of the fibroblast growth factor family. The complete mammalian FGF family. J Biol Chem. 2006;281:15694–700.

34. Gouaze-Andersson V, Delmas C, Taurand M, Martinez-Gala J, Evrard S, Mazoyer S, et al. FGFR1 Induces Glioblastoma Radioresistance through the PLCgamma/Hif1 alpha Pathway. Cancer research. 2016;76:3036–44.

35. Loilome W, Joshi AD, ap Rhys CM, Piccirillo S, Vescovi AL, Gallia GL, et al. Glioblastoma cell growth is suppressed by disruption of Fibroblast Growth Factor pathway signaling. Journal of neuro-oncology. 2009;94:359–66.

36. Morrison RS, Yamaguchi F, Saya H, Bruner JM, Yahanda AM, Donehower LA, et al. Basic fibroblast growth factor and fibroblast growth factor receptor I are implicated in the growth of human astrocytomas. Journal of neuro-oncology. 1994;18:207–16.

37. Gouaze-Andersson V, Gherardi MJ, Lemarie A, Gilhodes J, Lubrano V, Arnauduc F, et al. FGFR1/FOXM1 pathway: a key regulator of glioblastoma stem cells radioresistance and a prognosis biomarker. Oncotarget. 2018;9:31637–49.

38. Ohashi R, Matsuda Y, Ishiwata T, Naito Z. Downregulation of fibroblast growth factor receptor 2 and its isoforms correlates with a high proliferation rate and poor prognosis in high-grade glioma. Oncology reports. 2014;32:1163–9.

39. Toedt G, Barbus S, Wolter M, Felsberg J, Tews B, Blond F, et al. Molecular signatures classify astrocytic gliomas by IDH1 mutation status. International journal of cancer. 2011;128:1095–103.

40. Darmanis S, Sloan SA, Croote D, Mignardi M, Chernikova S, Samghababi P, et al. Single-Cell RNA-Seq Analysis of Infiltrating Neoplastic Cells at the Migrating Front of Human Glioblastoma. Cell Rep. 2017;21:1399–410.

41. Singh D, Chan JM, Zoppoli P, Niola F, Sullivan R, Castano A, et al. Transforming fusions of FGFR and TACC genes in human glioblastoma. Science. 2012;337:1231–5.

42. Rosmaninho P, Mukusch S, Piscopo V, Teixeira V, Raposo AA, Warta R, et al. Zeb1 potentiates genome-wide gene transcription with Lef1 to promote glioblastoma cell invasion. EMBO J. 2018;37.

43. Pearson A, Smyth E, Babina IS, Herrera-Abreu MT, Tarazona N, Peckitt C, et al. High-Level Clonal FGFR Amplification and Response to FGFR Inhibition in a Translational Clinical Trial. Cancer Discov. 2016;6:838–51.

44. Alvarado AG, Thiagarajan PS, Mulkearns-Hubert EE, Silver DJ, Hale JS, Alban TJ, et al. Glioblastoma Cancer Stem Cells Evade Innate Immune Suppression of Self-Renewal through Reduced TLR4 Expression. Cell Stem Cell. 2017;20:450–61 e4.

45. Hale JS, Otvos B, Sinyuk M, Alvarado AG, Hitomi M, Stoltz K, et al. Cancer stem cell-specific scavenger receptor CD36 drives glioblastoma progression. Stem Cells. 2014;32:1746–58.

46. Lathia JD, Gallagher J, Heddleston JM, Wang J, Eyler CE, Macswords J, et al. Integrin alpha 6 regulates glioblastoma stem cells. Cell Stem Cell. 2010;6:421–32.

47. Silver DJ, Siebzehnrubl FA, Schildts MJ, Yachnis AT, Smith GM, Smith AA, et al. Chondroitin Sulfate Proteoglycans Potently Inhibit Invasion and Serve as a Central Organizer of the Brain Tumor Microenvironment. The Journal of neuroscience: the official journal of the Society for Neuroscience. 2013;33:15603–17.

48. Kim JW, Botvinnik OB, Abudayyeh O, Birger C, Rosenbluh J, Shrestha Y, et al. Characterizing genomic alterations in cancer by complementary functional associations. Nature biotechnology. 2016;34:539–46.

49. Reich M, Liefeld T, Gould J, Lerner J, Tamayo P, Mesirov JP. GenePattern 2.0. Nature genetics. 2006;38:500–1.

50. Cerami E, Gao J, Dogrusoz U, Gross BE, Sumer SO, Aksoy BA, et al. The cBio cancer genomics portal: an open platform for exploring multidimensional cancer genomics data. Cancer Discov. 2012;2:401–4.

51. Subramanian A, Tamayo P, Mootha VK, Mukherjee S, Ebert BL, Gillette MA, et al. Gene set enrichment analysis: a knowledge-based approach for interpreting genome-wide expression profiles. Proceedings of the National Academy of Sciences of the United States of America. 2005;102:15545–50.

52. Liberzon A, Birger C, Thorvaldsdottir H, Ghandi M, Mesirov JP, Tamayo P. The Molecular Signatures Database (MSigDB) hallmark gene set collection. Cell systems. 2015;1:417–25.

