## Supplementary figures and images for "ADAMDEC1 maintains a novel growth factor signaling loop in cancer stem cells"

### Figure S1

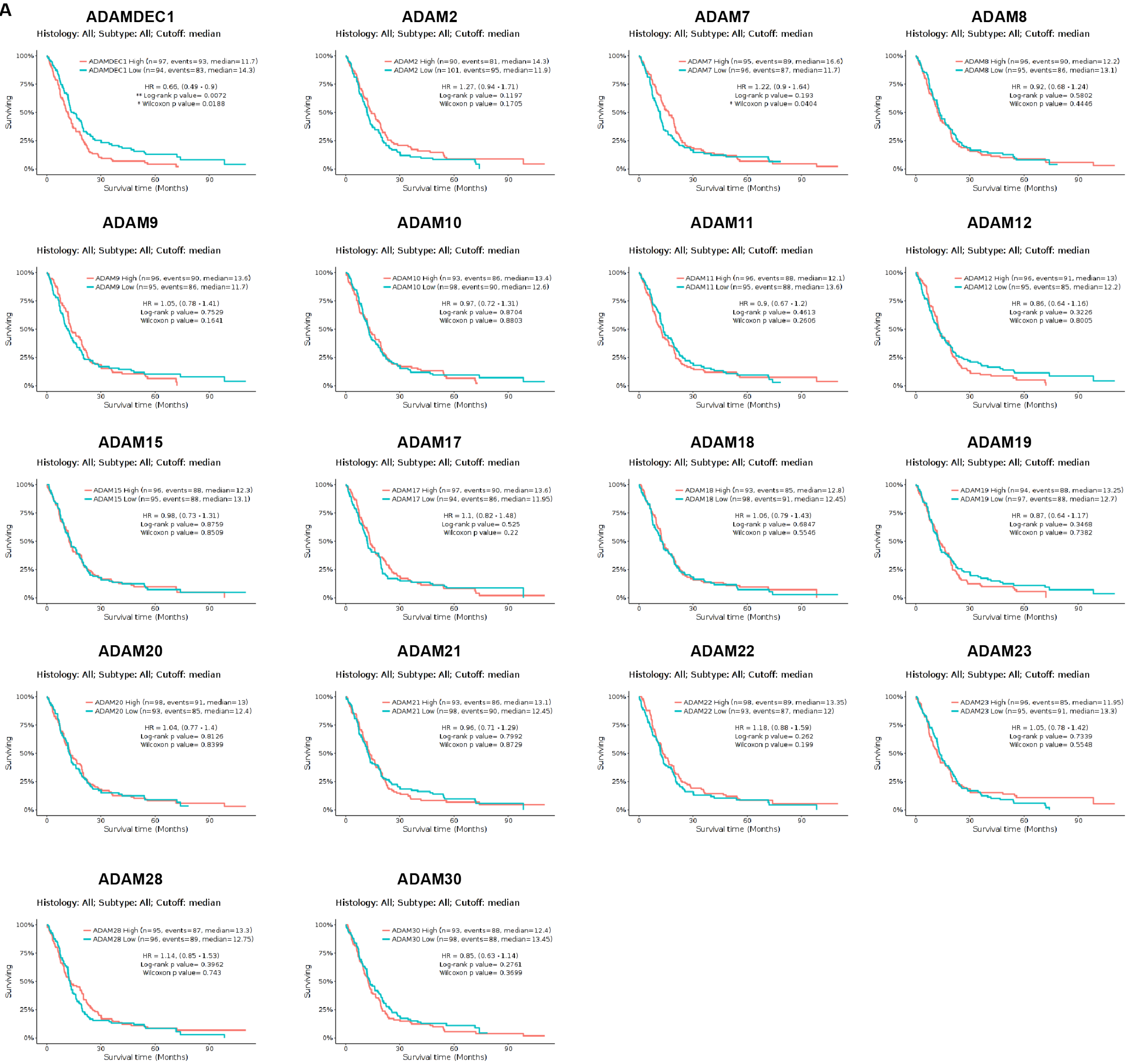

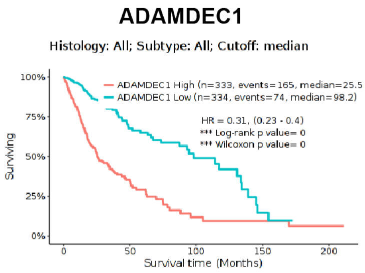

### Figure S2

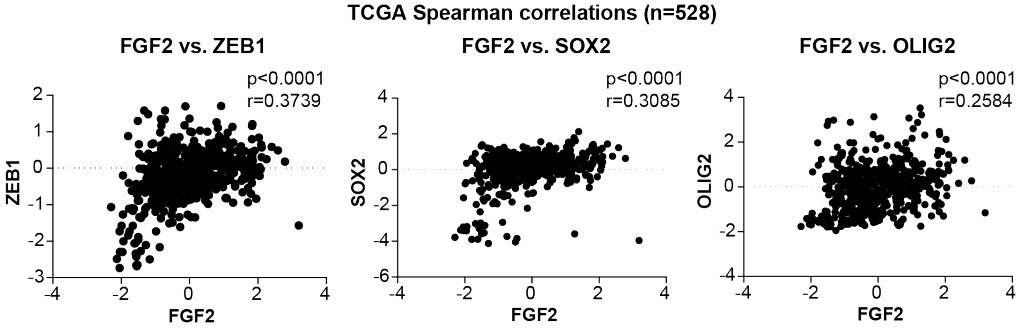

### Figure S3

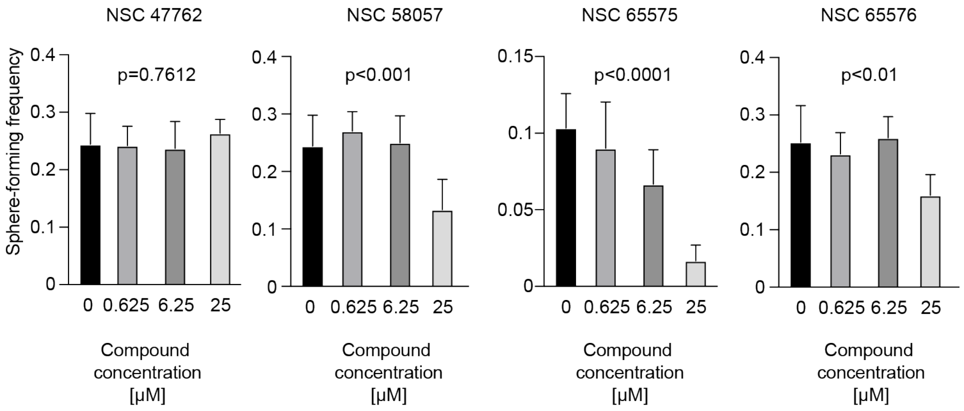

### Figure S4

#### Fig. S4

###### GSEA: Verhaak Glioblastoma Gene Sets (separate cluster vs all samples)

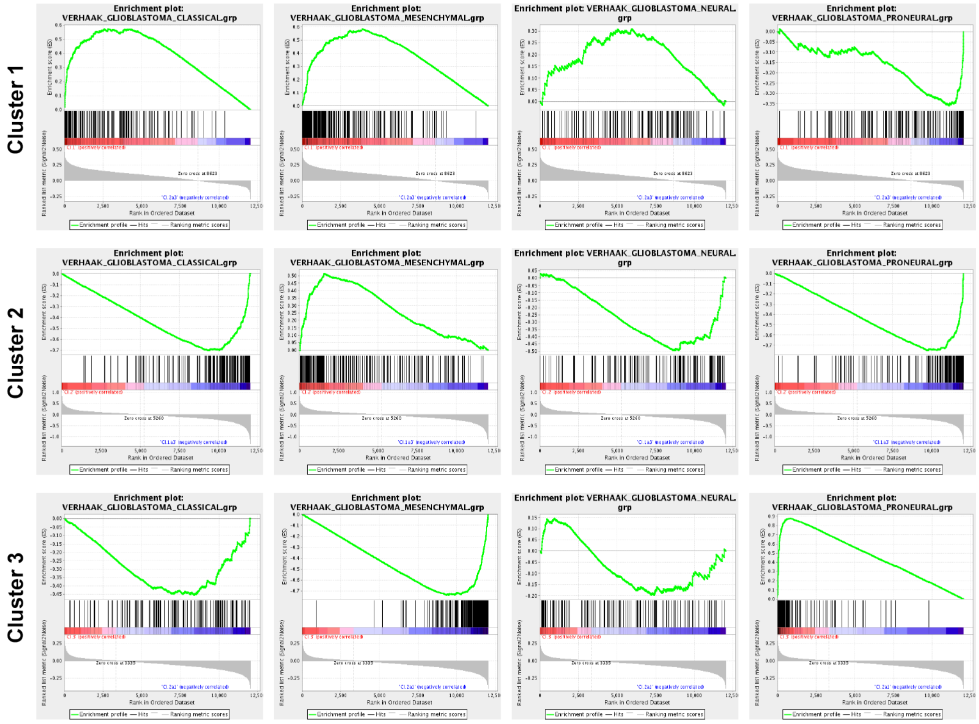

### Figure S5

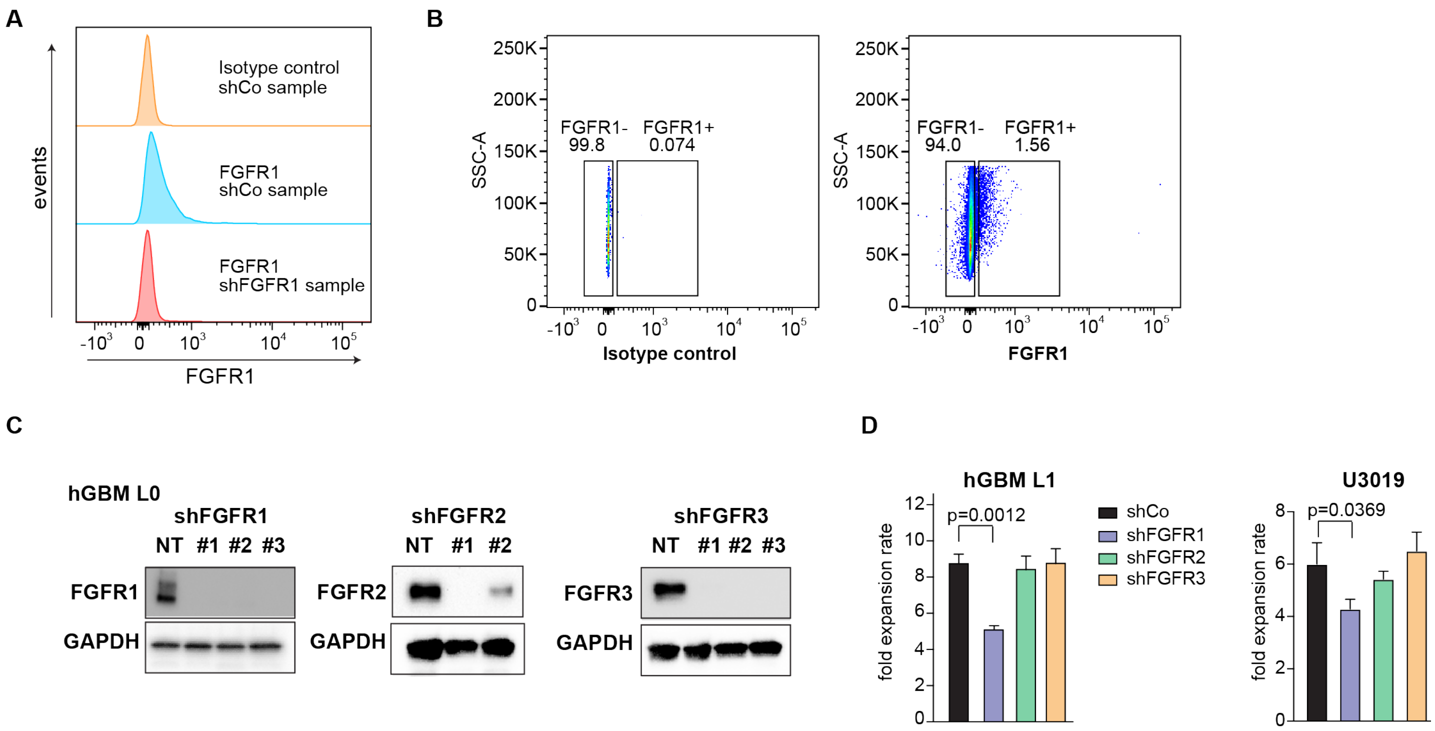

### Figure S6

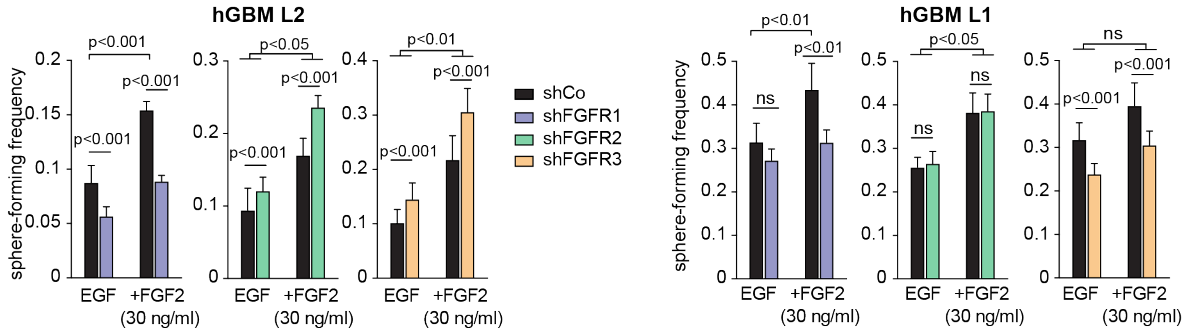

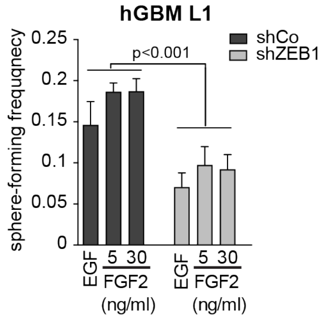
